# Concepts for structured illumination microscopy with extended axial resolution through mirrored illumination

**DOI:** 10.1101/828632

**Authors:** James D. Manton, Florian Ströhl, Reto Fiolka, Clemens F. Kaminski, Eric J. Rees

## Abstract

Wide-field fluorescence microscopy, while much faster than confocal microscopy, suffers from a lack of optical sectioning and poor axial resolution. 3D structured illumination microscopy (SIM) has been demonstrated to provide optical sectioning and to double the resolution limit both laterally and axially, but even with this the axial resolution is still worse than the lateral resolution of unmodified wide-field microscopy. Interferometric schemes using two high numerical aperture objectives, such as 4Pi confocal and I^5^M microscopy, have improved the axial resolution beyond that of the lateral, but at the cost of a significantly more complex optical setup. Here, we investigate a simpler dual-objective scheme which we propose can be easily added to an existing 3D-SIM microscope, providing lateral and axial resolutions in excess of 125 nm with conventional fluorophores and without the need for interferometric detection.

## 1 Introduction

Over the past three decades, there has been a considerable effort to improve the spatial resolution of fluorescence microscopy, with the majority of advances being focussed on improving the lateral resolution [1–16]. However, in many cases, improving the axial resolution would be just as, if not more, beneficial, particularly as the axial resolution of a conventional microscope is much worse than its lateral resolution. While both lateral and axial resolution increase with numerical aperture (NA), even an ideal single objective system that could collect over a full hemisphere on one side of the sample is bound to have an axial resolution at best half that of the lateral resolution. Such anisotropic resolution is highly detrimental for the accurate quantification of object sizes, shapes, volumes and curvatures.

3D structured illumination microscopy (3D-SIM) has been shown to double both lateral and axial resolution in wide-field microscopy through patterned illumination generated by the interference of three beams [17]. As well as developments in improving temporal resolution [18, 19], it has successfully been combined with dual-objective interferometric illumination and detection in I^5^S microscopy to achieve an unprecedented wide-field 3D isotropic resolution of 90 nm [20, 21]. Despite this impressive feat, use of this technology has been extremely limited due to the experimental difficulties in constructing and operating such a system. Even the use of the related dual-objective confocal technique of 4Pi microscopy has been limited by its complexity, with most existing systems relying on the simplest of the three 4Pi methods, Type A [22].

The main difficulty arising in implementing Type C 4Pi or I^5^S microscopy, in which both illumination and emission light are made to interfere, is ensuring that the path lengths from sample to beamsplitter in the interferometric arms are equal to within the coherence length of the fluorescence light (i.e. a few microns). In contrast, the relatively long coherence lengths of lasers commonly used in fluorescence microscopy ensures that interfering the excitation light, as in Type A 4Pi microscopy, is comparatively straightforward.

In this article, we briefly review the theory of 3D-SIM and use this to propose a simple dual-objective scheme that does not require interferometric detection to further improve the achievable axial resolution. In addition, by relaxing the technical constraints on the secondary objective, our scheme is compatible with low NA, high working distance objectives, further simplifying an experimental realisation.

We first review the theory of operation of 3D-SIM and use this to introduce our method. We present an analysis of the potential performance of our approach via geometric considerations and support this with numerical simulations of the image formation process, providing examples of expected images given a known ground truth object. We detail the practical considerations for an experimental realisation of our method and propose that existing 3D-SIM microscopes can be easily modified to further double axial resolution while maintaining the same reconstruction approach.

### 1.1 3D-SIM theory

Considering a fluorescence microscope as a linear, shift-invariant imaging system, the image data collected, *D*, can be considered as the convolution of the fluorescence emission, *F*, with a point spread function (PSF), *H*, such that *D* = *F* * *H* = (*S* × *I*) * *H*, where *S* is the sample fluorophore distribution and *I* is the illumination intensity distribution. Alternatively, in the Fourier domain, 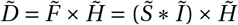, where overset tildes denote the Fourier transforms of the respective real-space functions and 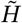 is the optical transfer function (OTF) [5–7].

The illumination intensity, *I*, is related to the electric field, *E*, via *I* = *EE*^*^, where *E*^*^ is the complex conjugate of *E*. Hence, the Fourier domain distribution, *Ĩ*, is given by the autocorrelation of 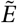. In a conventional 3D-SIM system, the illumination is generated by three interfering laser beams produced by the 0 and ±1 diffraction orders of a grating. This means that the Fourier domain electric field amplitude distribution, 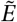, consists of three points on a shell of radius 2*π*/*λ*, spaced angularly by no more than *α*, where *λ* is the wavelength of the illumination light (given by *λ*_0_/*n*, where *λ*_0_ is the vacuum wavelength and *n* is the refractive index of the medium) and *α* is the half-angle of the objective lens (shown in magenta in Figure 1a). The Fourier domain distribution, *Ĩ*, hence consists of the seven magenta points shown in Figure 1c [17].

The detection system of the microscope cannot capture information outside the support of the OTF, but this illumination structure down-mixes higher spatial frequency information into the passband of the detection objective. Under the assumption that the illumination pattern remains fixed with respect to the objective as *z*-stacks are collected, it has previously been shown by Gustafsson et al. that the components are already unmixed axially by the process of moving the sample through focus [17]. All that remains is to unmix laterally, by applying lateral phase shifts to the illumination pattern, producing as manybands ofinformation as there are columns of components in the illumination Fourier domain distribution. In this case, while there are seven components in total (magenta points in Figure 1c), only five phase steps are required. By repeating this process for multiple orientations of the illumination pattern, the lateral resolution enhancement is made approximately isotropic. Typically three orientations are used, requiring 15 images per focal plane. We can consider the effective overall OTF as the combination of all the axially extended bands produced by this process, shown with a magenta fill in Figure 1c.

## 2 Results

### 2.1 Further improving axial resolution in SIM by including a fourth illumination beam

Consider a modified 3D-SIM system in which a further centralbeamisincidentuponthe sample fromthe opposite side. Here, the Fourier domain electric field amplitude distribution includes an additional point on the shell of radius 2*π*/*λ* opposite the original central beam (green point in Figure 1a). This causes the Fourier domain illumination intensity distribution to gain a further six points, drawn in green in Figure 1c.

As before, we can describe the effective overall OTF as the combination of all the axially extended bands produced by this process, the bandlimit of which shown as the black line in Figure 1c. Here, six copies of the wide-field OTF (shown in green) have been added above and below the original 3D-SIM OTF, providing enhanced optical sectioning and axial resolution. Because the additional beam is incident on-axis, no extra lateral frequency components are created. This means that the lateral unmixing process still requires only five phase shifts, but as the axial support of the OTF is enlarged *z*-stacks must be acquired with finer steps in order to satisfy the Nyquist sampling criterion.

### 2.2 Geometric considerations for filling the holes in the OTF

Taking the 13-point illumination structure and convolving with the conventional wide-field OTF produces a system OTF with larger axial support but which, crucially, contains holes (white fill in Figure 1c). While considerably smaller than those present in standing-wave microscopy, in order for all sample frequencies within the maximum extent of the OTF to be accurately recorded these holes must be filled [23]. This cannot be achieved by merely changing the angular spread of the outer points of the illumination and so alternative approaches must be sought. Here, we consider three approaches to achieve this. In the following discussion, examples are shown for an oil-immersion 1.45 NA lens operating in a nominal refractive index of 1.518. Unless otherwise stated, illumination is at 488 nm with fluorescence emission detected at 510 nm.

#### 2.2.1 Illuminating with multiple wavelengths

Consider the effect of illuminating with two different wavelengths. While the overall structure of the illumination will be similar in each case, the wavelength difference will cause the distance from the origin of each Fourier domain illumination component to differ (compare purple and green points in Figure 2a). By selecting an appropriate combination of wavelengths, such as 445 nm and 561 nm illumination with 610 nm detection, the holes for each wavelength can be filled with information obtained using the other illumination wavelength. With this, nine phase steps rather than five are required (as there are nine distinct columns of components), slowing acquisition speed. In addition, the structure of interest now needs to be labelled with fluorophores with a broad excitation spectrum, such as quantum dots, complicating multicolour imaging of different structures. Alternatively, two different fluorophores with well-separated excitation and emission spectra can be used, with two sets of five-phase data acquired and fused, at the expense of a further reduction in acquisition speed.

#### 2.2.2 Illuminating with multiple modes

Instead of illuminating with multiple wavelengths, consider illuminating not with single modes in the pupil, but with multiple independent incoherent modes over a significant area of the pupil. Each order generated by the grating produces an image of the source in the pupil. For appropriately spatially incoherent illumination, each point within each source image will be incoherent with all other points within the same image, but coherent with the corresponding points in the other source images [17].

As the objective lens obeys the Abbe sine condition, points in the pupil are projected orthographically onto the shell of radius 2*π*/*λ* when considering the Fourier domain electric field amplitude distribution. Hence, for each set of source point images, the lateral components of the Fourier domain illumination intensity distribution will not change while the axial ones will. The end effect, after summing over all points within the source images, is an illumination intensity distribution in which each of the original points (except for ones along *k_z_*) have been broadened axially but not laterally (green and magenta lines in Figure 2c).

**Figure 1:**
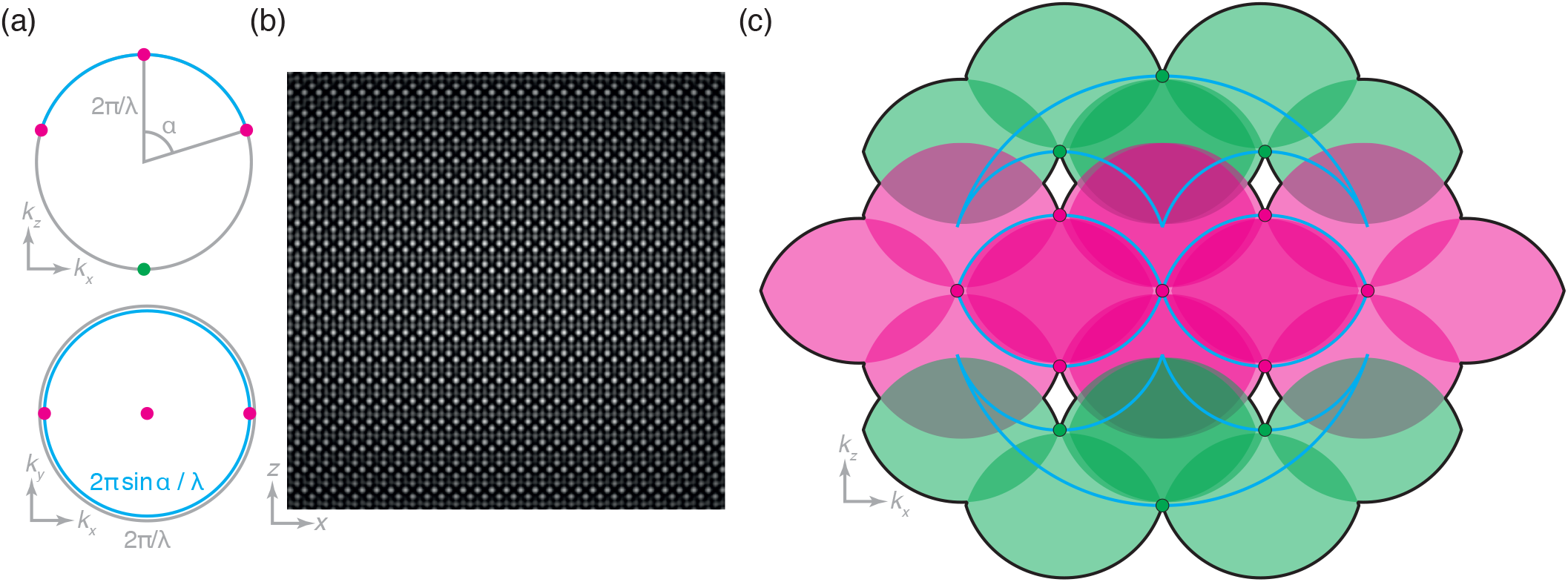
Creating an illumination profile using four mutually coherent beams. (a) Fourier domain illumination electric field distribution formed by four coherent beams, with those present in 3D-SIM drawn in magenta and the additional beam considered in this work drawn in green. Here the angular separation is the maximum possible, namely the half-angle, *α*, of the lens. (b) Real domain illumination intensity distribution resulting from the electric field distribution of (a). (c) Fourier domain illumination intensity distribution drawn as magenta and green spots (corresponding to those present in 3D-SIM and the new components considered in this work) with the overall support of the OTF drawn as a black line and each wide-field OTF contribution shown in the colour of the illumination component that produced it. These illumination components can be seen to lie on the edge of the support of the 4Pi OTF (cyan lines) for this case where the angular separation is *α*. Note the holes within the support of the OTF near the off-axis green spots.

The outermost components are broadened to the full axial extent of the wide-field OTF at that lateral frequency, while the inner components are broadened half as much. For a pupil fraction of 1/3 (the maximum possible before source images overlap), corresponding to a maximum lateral illumination frequency of 2/3 that of the maximum supported by the lens, the effect is such that all components are broadened to the full extent of the local component of the dual-objective OTF (compare green and magenta lines with cyan dual-objective OTF extent in Figure 2c).

While the exact absolute extent of the axial broadening and overlap is dependent on the exact numerical aperture, refractive index of immersion, *n*, and pupil fraction used, a pupil fraction of 1/3 is more than sufficient to fill all the holes for a 1.45 numerical aperture oil immersion objective (*n* = 1.518) as demonstrated in Figure 2c. While this approach does not require any changes to the sample labelling and fills the holes with single-orientation reconstruction procedures, the maximum possible lateral resolution enhancement is decreased as the illumination pattern is formed by beams spaced by a lower numerical aperture.

#### 2.2.3 Recording data with multiple pattern orientations

So far we have only considered a two-dimensional system operating in the *xz*-plane (where *z* is the optical axis). In a true three-dimensional system the wide-field OTF is not the bi-lobed object depicted in cyan in Figure 1c, but the toroidal solid of revolution given by that cross-section [24]. Hence, by recording data with three pattern orientations spaced by 120°, the holes in one orientation are filled by contributions from the others (as shown in Figure 2b). Therefore, illuminating with multiple wavelength or with multiple modes is not strictly necessary if the numerical aperture of the detection objective is sufficiently large. However, this precludes acquisitions using just one orientation, which may be desired for reasons of speed when enhancing the axial resolution is more important than an isotropic lateral resolution enhancement.

### 2.3 Numerical simulations of imaging performance

In order to validate our conclusions based on a geometrical understanding of the optical transfer functions relevant to our proposed approach, we produced numerical simulations of imaging performance based on a scalar, non-paraxial model of diffraction. Figure 3 shows simulated modulation transfer functions for conventional wide-field microscopy, I^5^M [25], 3D-SIM, our proposed approach, six-beam structured illumination microscopy (i.e. I^5^S without interferometric detection), and I^5^S microscopy.

Detection OTFs were produced by autocorrelating caps of the Ewald sphere, assuming an aperture half-angle of 72.7° (corresponding to a 1.27 NA water immersion objective or a 1.45 oil immersion objective for index-matched samples). The illumination intensity Fourier distribution was produced by autocorrelating appropriate sections of the Ewald sphere, with the result being Fourier transformed to produce a real space illumination function. For structured illumination techniques, the ±1 orders were set to lie at 95 % of the pupil radius, with three orientations of the pattern used to generate the overall illumination function. The overall OTF was produced by multiplying the detection point spread function (produced by Fourier transforming the calculated OTF) with the illumination function and Fourier transforming the result.

**Figure 2:**
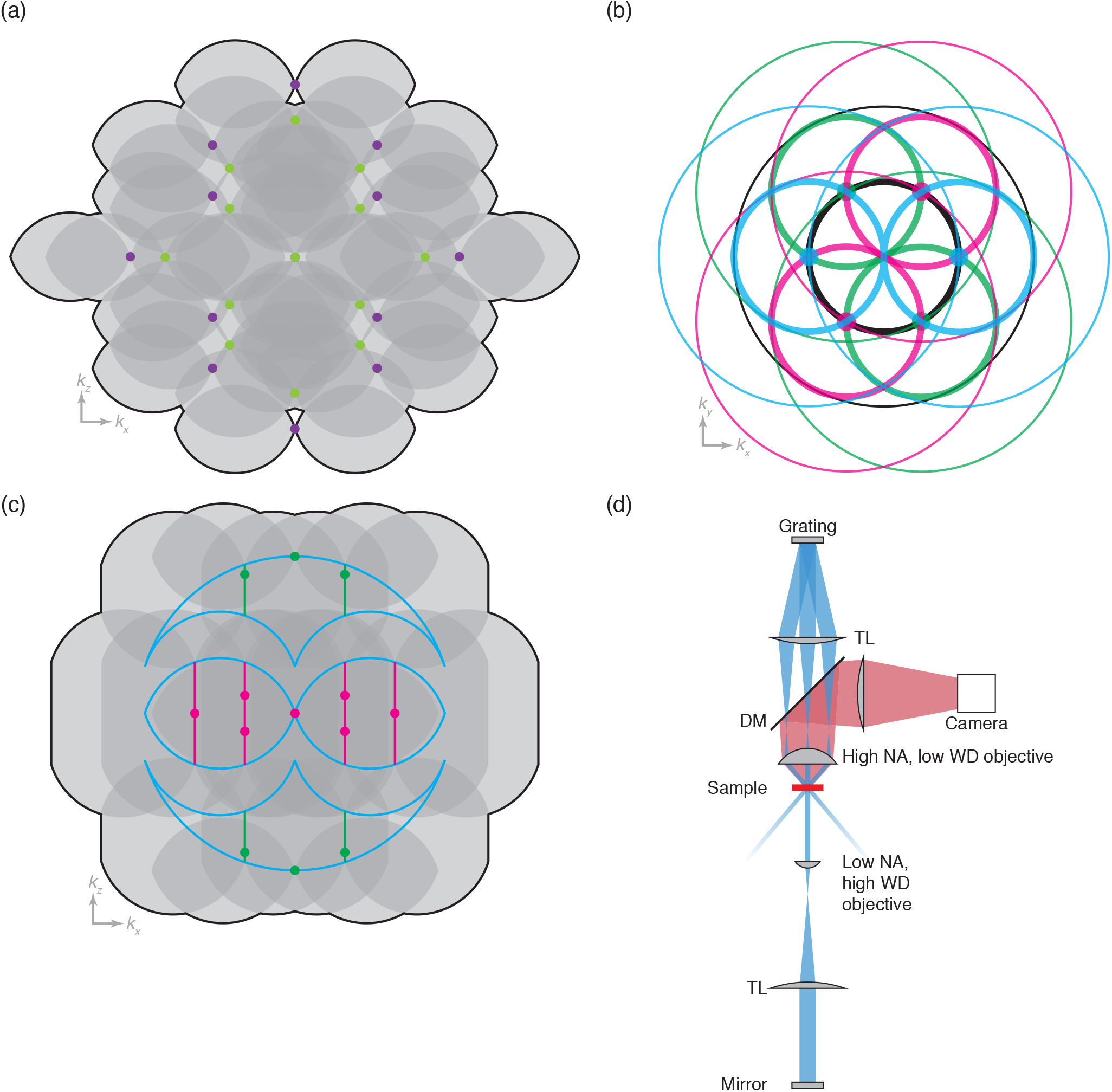
Strategies for filling the holes in the OTF. (a) Overall effective OTF (support shown as black line) for illumination at 445 nm (purple spots) and 561 nm (yellow-green spots) with fluorescence detection at 610 nm (gray OTFs). Here the holes are filled due to the difference in *k*-vector length, but two new ones are created along *k_z_*. These can easily be removed by apodisation at the cost of slightly reduced resolution. (b) The effect of considering multiple orientations on the *k_y_*–*k_x_* coverage. The ever-present zero order wide-field detection OTF component is shown in black, with the thin line denoting the edge of the OTF support, where the axial extent is lowest, and the thick black line denoting the region of the OTF where the axial extent is highest. The first order contributions from each orientation are shown in cyan, magenta and green, with the holes in the OTF support from each orientation being shown by the large filled circles. As can be seen from the overlap of thick lines with circles, each hole is effectively filled in by contributions from the next orientation. (c) The effect of multimode illumination with a pupil fraction of 1/3. Considering the centre of each illumination image created by the grating leads to the Fourier domain illumination distribution shown as the magenta and green spots. Summing over all such point tetrads causes an axial broadening of this distribution, shown as the magenta and green lines. While no contributions from outside the 4Pi OTF exist, the shifting of contributions within this ensures a good overlap and hence no holes in the overall OTF support. (d) Schematic of the proposed experimental setup in which a conventional 3D-SIM microscope is augmented with a low numerical aperture, high working distance objective on the other side of the sample, paired with a tube lens and mirror to reflect just the central beam.

In the displayed results, OTFs labelled ‘slice’ correspond to a slice of the OTF in the Fourier domain, not a slice of the image data in the real domain. Similarly, OTFs labelled ‘projection’ correspond to a sum of the OTF along the unshown axis (i.e. orthogonal to the page) and so, by the projection-slice theorem, correspond to a slice of the image data in the real domain (i.e. these are the equivalent to the result of Fourier transforming the image data corresponding to a slice of a sub-diffraction point source). Code to reproduce these results is presented in Supplementary Code File 1.

The OTFs shown in Figure 3d support our conclusion in Section 2.2.3 that multiple orientations alone are sufficient to fill the holes in the OTF, without the need for illumination at two wavelengths (Section 2.2.1) or with multiple modes (Section 2.2.2). Indeed, here the improvements in optical sectioning and axial resolution are clearly shown by the high value of the transfer function along *k_z_* away from the origin — only at the very edge of the support does the value fall below that shown in the corresponding lateral transfer function at the usual diffraction limit in the 3D-SIM case (see Figure 3c). Supplementary Video 1 shows the effect of increasing the primary objective NA on the OTF and clearly shows that the holes are well-filled by a 1.2 NA water immersion lens(or, equivalently, a 1.37 NA oil immersion lens). Experimentally, OTFs are always found to be weaker than theory suggests, supporting the idea that the holes should be well-filled in our simulations in order for them to be filled in practice. In addition, we see that the holes are not filled at an NA of 1.1, the highest commercially available NA for awater-dipping objective.

To more readily appreciate the expected improvement in optical sectioning and axial resolution given by our technique over 3D-SIM, we used these OTFs to produce simulated images given a constructed ground truth (see Figure 4). Here, an *xz* slice of an artificial ground truth structure is shown in Figure 4a. Corresponding image data slices for 3D-SIM, I^5^S, and our proposed four-beam SIM method are shown in Figure 4b–d. Here, we have purposefully neglected to include the effects of noise to more clearly show the effects of the respective transfer functions.

First, we note that the I^5^S image shows the best contrast, as expected from its more uniform transfer function. Second, we note that both 3D-SIM and our proposed method have similar contrast, but that our method clearly separates closely-spaced fibrilar structures that are resolved as one thicker object in the 3D-SIM case (cyan arrowheads in Figure 4). Finally, while the resolution of I^5^S is theoretically superior to our proposed method due to its larger axial support, it is hard to appreciate this in the raw image data, suggesting that our method may provide almost all of the practical resolution improvements of I^5^S despite its relative simplicity.

### 2.4 Considerations for an experimental realisation

The required fourth beam can be provided via a relatively simple modification to an existing 3D-SIM system. By placing a low numerical aperture, high working distance objective on the opposite side of the sample, the central beam can be captured while the outer beams are left to escape. A tube lens and mirror placed in the image plane reflects this central beam, sending a fourth beam into the sample as required (see Figure 2d).

As we require this reflected beam to interfere with the three incident from the high NA objective side of the sample, the coherence length of the laser must be greater than the ~800 mm extra path length experienced by the reflected beam. Hence, for a 488 nm laser line, the frequency bandwidth must be less than ~100 MHz. While laser sources conventionally used for SIM may not have a sufficiently small bandwidth, appropriate single frequency lasers are readily available from commercial sources.

In order for the central beam to be fully captured by the low NA objective, the NA of this lens must be greater than the product of the NA of the high NA objective with the illumination pupil fraction. Objective lenses with NAs of 0.5 and working distances greater than 1mmare commercially available and would be suitable for capturing the central beam from a 1.45 NA primary objective, even if the pupil fraction is the maximum possible (1/3).

If the multimode approach is to be used, illumination can be provided by laser light coupled into a shaken multimode fibre as this well-approximates a spatially incoherent source over sufficiently long times (typically of the order of less than a millisecond) [19]. As the diameter of the objective pupil is given by 2*f* × NA, where *f* is the back focal length and NA is the numerical aperture, a 1/3 pupil fraction would correspond to a ~20-fold magnification for a 100 μm core fibre and a 100× objective designed for a 200 mm tube length. Such fibres are commonly available with a numerical aperture of 0.22, ensuring minimal loss of light through the relay optics from fibre tip to pupil image while still satisfying the conditions for proper demagnification of the grating onto the sample.

Practically, only a small pupil fraction would be necessary, with the multimode illumination being used mainly to reduce laser speckle and other coherence artefacts while the multiple pattern orientations fill the holes in the OTF. With this in mind, we can consider using a water-dipping lens as the secondary objective. This would make our method compatible with imaging samples loaded into coverglass-bottomed wells, with easy access to the sample for electrophysiology, treatment delivery and media exchange maintained. This is in contrast to the existing 4Pi confocal, I^5^M and I^5^S methods which require the sample to be mounted within a coverglass sandwich as they require high-NA immersion lenses.

## 3 Discussion

We have investigated the possibility of further axial resolution enhancement in 3D-SIM by reflecting the central beam and have provided methods for countering the existance of holes in the OTF support. To our surprise, we found that a fully three-dimensional consideration of the image formation process in structured illumination microscopy showed that these holes are well-filled when a full set of image data for all (three) pattern rotations are acquired and combined. Numerical simulations demonstrated a clear improvement in axial resolution over 3D-SIM, with a practical effective resolution close to that of I^5^S.

**Figure 3:**
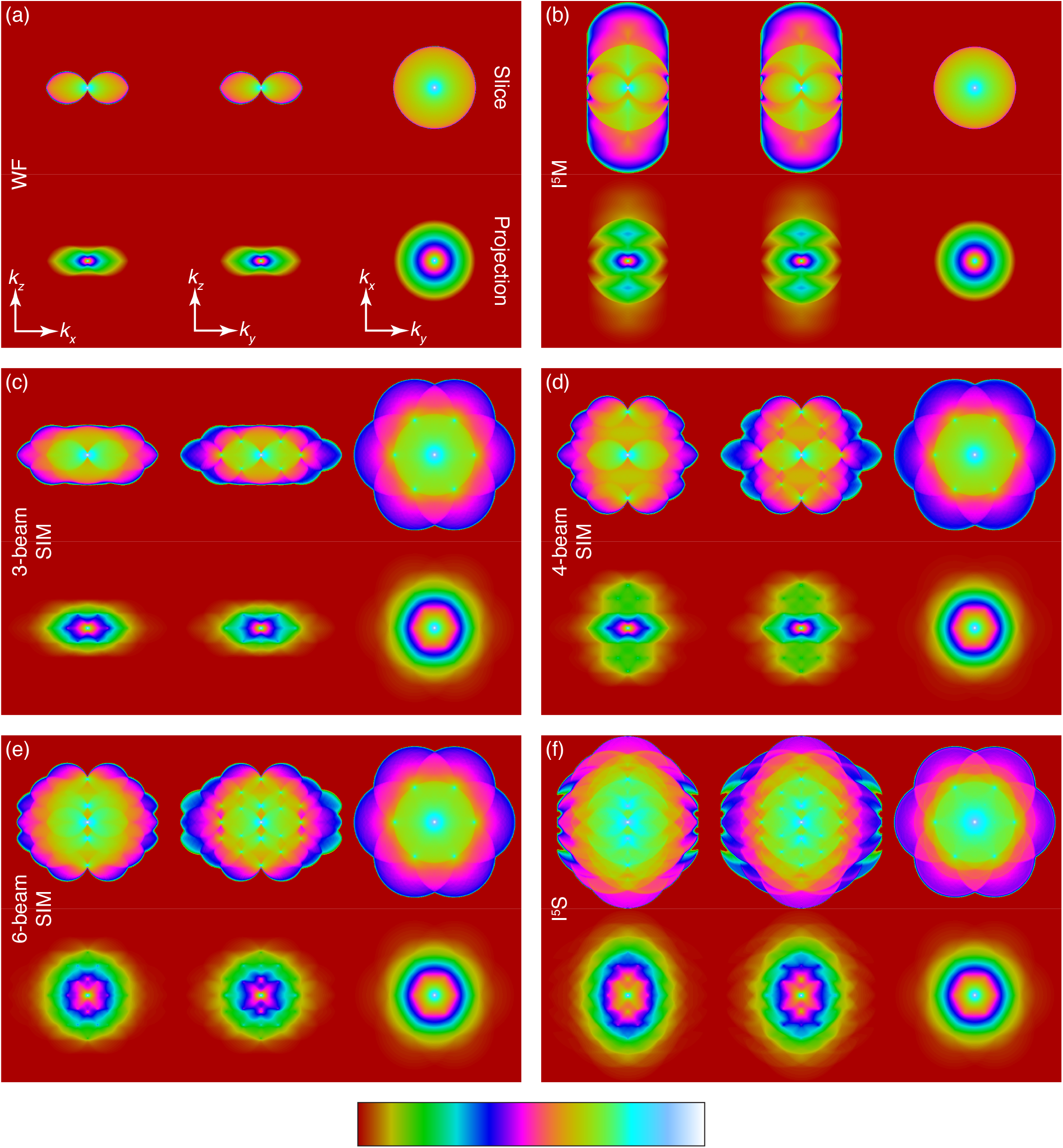
Simulated modulation transfer functions (MTF) for wide-field microscopy techniques, assuming an aperture half-angle of 72.7°. Top rows show the central slice of the MTF, while bottom rows show the projection of the MTF (as would be seen by Fourier transforming a slice of a sub-diffraction bead). All panels follow the same co-ordinate systems. Central slices are shown on a logarithmic scale to enhance dim details. (a) Wide-field microscopy with uniform illumination. (b) I^5^M (incoherent illumination + interferometric detection). (c) 3D-SIM. (d) Our proposed dual-objective four-beam SIM (without interferometric detection). (e) Dual-objective six-beam SIM without interferometric detection. (f) I^5^S (dual-objective six-beam SIM + interferometric detection).

**Figure 4:**
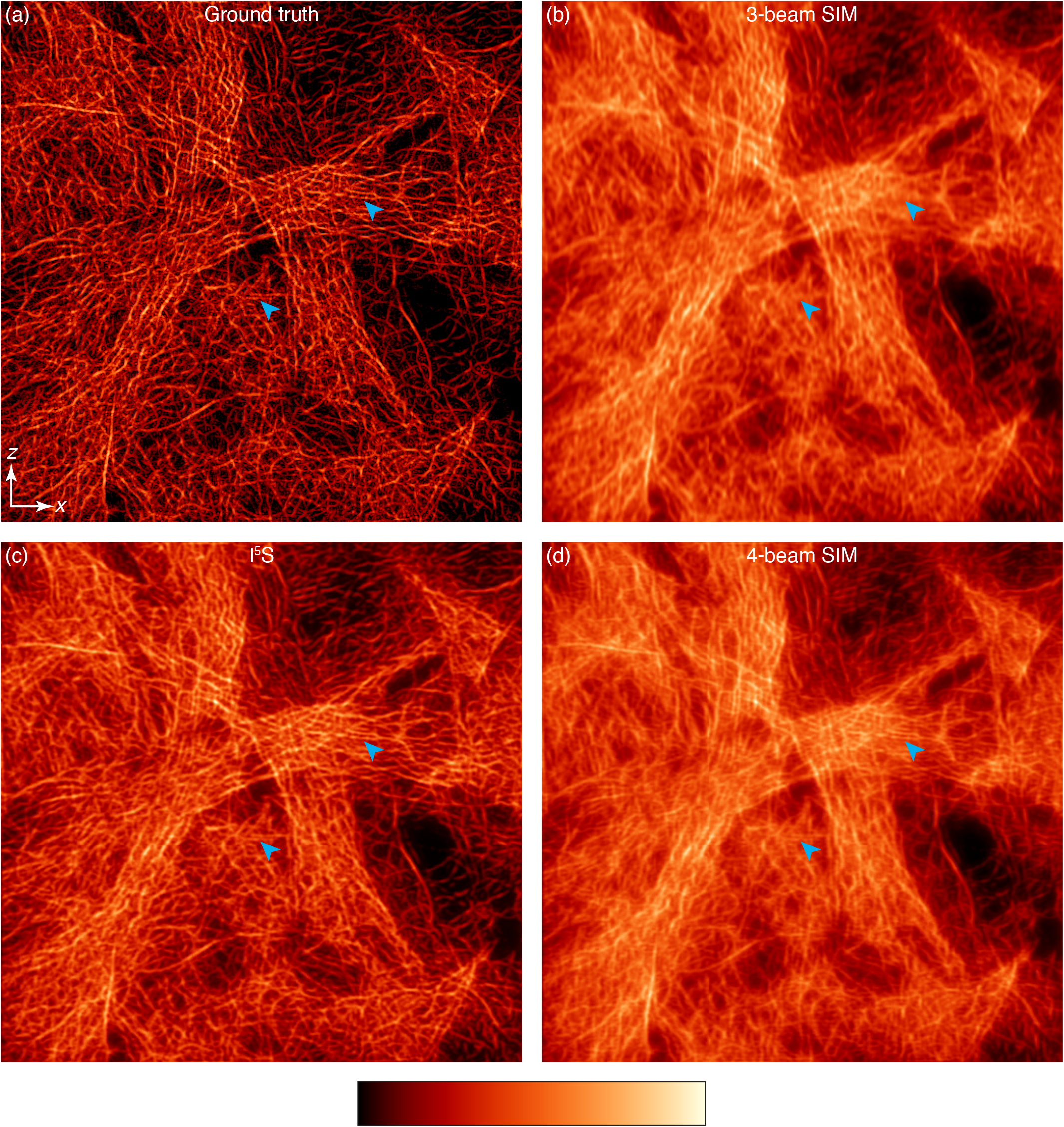
Simulated imaging performance on a fibrous ground truth test image, shown as an *xz* slice. All images are normalised to have the same minimum and maximum intensity. Cyan arrowheads point to closely-spaced fibre pairs that cannot be resolved using conventional 3D-SIM, but which are resolvable using our four-beam approach or I^5^S. (a) Ground truth. (b) 3D-SIM. (c) I^5^S (dual-objective six-beam SIM + interferometric detection). (d) Our proposed dual-objective four-beam SIM (without interferometric detection).

For an objective with an aperture half-angle of 72.7° (corresponding to a 1.27 NA water immersion objective or a 1.45 oil immersion objective for index-matched samples) a fully isotropic 3D resolution (as defined by OTF support) at ~1.5 × the diffraction-limited lateral resolution can be achieved by apodising a fully reconstructed four-beam SIM dataset using three pattern orientations. Ignoring a further resolution enhancement provided by the Stokes’ shift, this would convert a live-imaging system at 510 nm from a 200 nm × 200 nm × 545 nm resolution to a 135 nm × 135 nm × 135 nm resolution, i.e. an almost nine-fold increase in volumetric information density.

While further resolution enhancements would be possible by using another high NA lens as the secondary objective to provide a further two illumination beams, and/or by using interferometric detection as in I^5^M and I^5^S microscopy [20, 25], the comparatively simple experimental setup proposed here achieves much of the performance of such a system while avoiding the significant complexity of interferometric detection. By relaxing the requirements of the secondary objective, our approach is compatible with imaging samples mounted in a conventional manner in multi-well plates, rather than sandwiched between two coverslips — a long-working-distance dipping lens is a sufficient secondary objective.

We emphasise that our approach is compatible with existing methods for reconstructing three-dimensional structured illumination microscopy data, with a minor modification to the shape of the OTF bands used. These are readily calculated or measured using exactly the same approaches as used for conventional 3D-SIM and so should be straightforward to integrate into existing sofware, such as fairSIM [26].

## Supplementary Information

**Supplementary Video 1:**
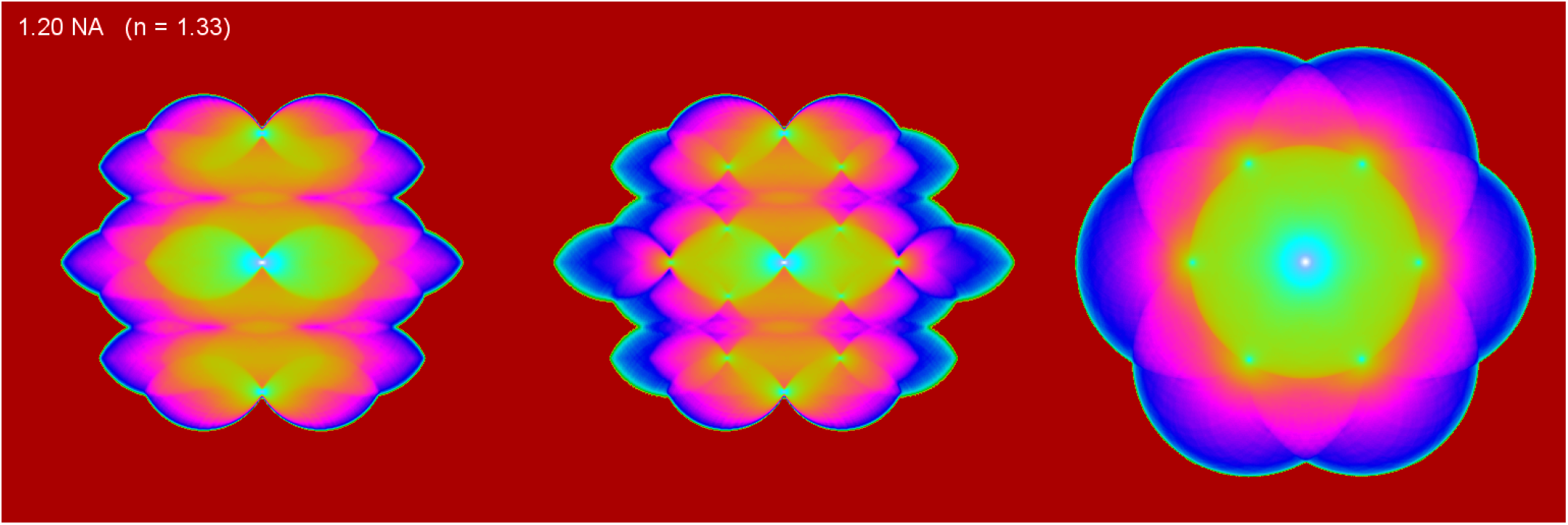
Simulated optical transfer functions for our four-beam SIM approach showing the effect of increasing primary objective numerical aperture (NA). Central slices are shown, displayed using the same co-ordinate system, logarithmic scaling and colourmap as in Figure 3. Holes are shown to be well-filled by an NA of 1.2, corresponding to commonly available water-immersion objectives.

**Supplementary Video 2:**
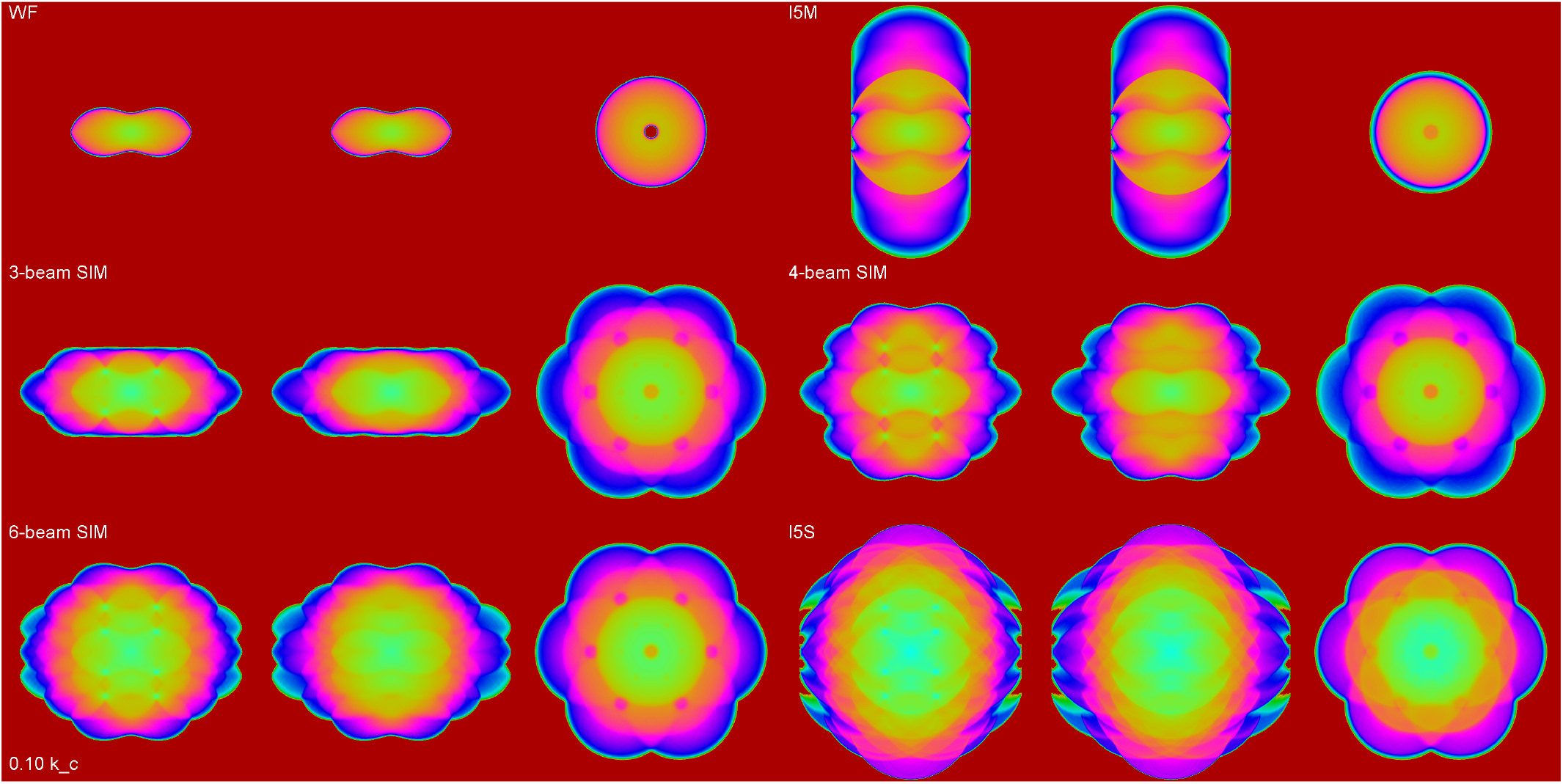
Simulated optical transfer functions for the microscopy techniques shown in Figure 3, showing orthogonal cuts, assuming a 72.7° aperture half-angle (1.27 NA water immersion, 1.4 NA oil immersion). Displays use the same co-ordinate system, logarithmic scaling and colourmap as in Figure 3. *k_c_* refers to the critical wavenumber, given by 2*πn*/4*λ*, where *n* is the refractive index.

## References

1. C. J. R. Sheppard. (1988). “Super-resolution in Confocal Imaging”. Optik 80, 83–84.

2. Stefan W. Hell and Jan Wichmann. (1994). “Breaking the diffraction resolution limit by stimulated emission: stimulated-emission-depletion fluorescence microscopy”. Optics Letters 19, 780. doi: 10.1364/OL.19.000780.

3. Stefan W. Hell. (1994). “Improvement of lateral resolution in far-field fluorescence light microscopy by using two-photon excitation with offset beams”. Optics Communications 106, 19–24. doi: 10.1016/0030-4018(94)90050-7.

4. E. Betzig. (1995). “Proposed method for molecular optical imaging”. Optics Letters 20, 237. doi: 10.1364/OL.20.000237.

5. Rainer Heintzmann and Christoph G. Cremer. (1999). “Laterally modulated excitation microscopy: improvement of resolution by using a diffraction grating”. BiOS Europe’98, 185–196. doi: 10.1117/12.336833.

6. M. G. L. Gustafsson. (2000). “Surpassing the lateral resolution limit by a factor of two using structured illumination microscopy”. Journal of Microscopy 198, 82–87. doi: 10.1046/j.1365-2818.2000.00710.x.

7. Jan T. Frohn, Helmut F. Knapp, and Andreas Stemmer. (2000). “True optical resolution beyond the Rayleigh limit achieved by standing wave illumination”. Proceedings of the National Academy of Sciences 97, 7232–7236. doi: 10.1073/pnas.130181797.

8. George E. Cragg and Peter T. C. So. (2000). “Lateral resolution enhancement with standing evanescent waves”. Optics Letters 25, 46. doi: 10.1364/OL.25.000046.

9. Rainer Heintzmann. (2003). “Saturated patterned excitation microscopy with two-dimensional excitation patterns”. Micron 34, 283–291. doi: 10.1016/S0968-4328(03)00053-2.

10. Mats G. L. Gustafsson. (2005). “Nonlinear structured-illumination microscopy: Wide-field fluorescence imaging with theoretically unlimited resolution”. Proceedings of the National Academy of Sciences of the United States of America 102, 13081–13086. doi: 10.1073/pnas.0406877102.

11. Keith A. Lidke et al. (2005). “Superresolution by localization of quantum dots using blinking statistics”. Optics Express 13, 7052–7062. doi: 10.1364/OPEX.13.007052.

12. Eric Betzig et al. (2006). “Imaging Intracellular Fluorescent Proteins at Nanometer Resolution”. Science 313, 1642–1645. doi: 10.1126/science.1127344.

13. Samuel T. Hess, Thanu P. K. Girirajan, and Michael D. Mason. (2006). “Ultra-High Resolution Imaging by Fluorescence Photoactivation Localization Microscopy”. Biophysical Journal 91, 4258–4272. doi: 10.1529/biophysj.106.091116.

14. Michael J. Rust, Mark Bates, and Xiaowei Zhuang. (2006). “Sub-diffraction-limit imaging by stochastic optical reconstruction microscopy (STORM)”. Nature Methods 3, 793–796. doi: 10.1038/nmeth929.

15. Alexey Sharonov and Robin M. Hochstrasser. (2006). “Wide-field subdiffraction imaging by accumulated binding of diffusing probes”. Proceedings of the National Academy of Sciences 103, 18911–18916. doi: 10.1073/pnas.0609643104.

16. Claus B. Müller and Jörg Enderlein. (2010). “Image Scanning Microscopy”. Physical Review Letters 104, 198101. doi: 10.1103/PhysRevLett.104.198101.

17. Mats G. L. Gustafsson et al. (2008). “Three-Dimensional Resolution Doubling in Wide-Field Fluorescence Microscopy by Structured Illumination”. Biophysical Journal 94, 4957–4970. doi: 10.1529/biophysj.107.120345.

18. Lin Shao et al. (2011). “Super-resolution 3D microscopy of live whole cells using structured illumination”. Nature Methods 8, 1044–1046. doi: 10.1038/nmeth.1734.

19. R. Fiolka et al. (2012). “Time-lapse two-color 3D imaging of live cells with doubled resolution using structured illumination”. Proceedings of the National Academy of Sciences 109, 5311–5315. doi: 10.1073/pnas.1119262109.

20. Lin Shao et al. (2008). “I5S: Wide-Field Light Microscopy with 100-nm-Scale Resolution in Three Dimensions”. Biophysical Journal 94, 4971–4983. doi: 10.1529/biophysj.107.120352.

21. L. Shao et al. (2012). “Interferometer-based structured-illumination microscopy utilizing complementary phase relationship through constructive and destructive image detection by two cameras”. Journal of Microscopy 246, 229–236. doi: 10.1111/j.1365-2818.2012.03604.x.

22. Stefan Hell and Ernst H. K. Stelzer. (1992). “Properties of a 4Pi confocal fluorescence microscope”. Journal of the Optical Society of America A 9, 2159. doi: 10.1364/JOSAA.9.002159.

23. Brent Bailey et al. (1993). “Enhancement of axial resolution in fluorescence microscopy by standing-wave excitation”. Nature 366, 44–48. doi: 10.1038/366044a0.

24. B. Roy Frieden. (1967). “Optical Transfer of the Three-Dimensional Object*†”. JOSA 57, 56–66. doi: 10.1364/JOSA.57.000056.

25. M. G. Gustafsson, D. A. Agard, and J. W. Sedat. (1999). “I5M: 3D widefield light microscopy with better than 100 nm axial resolution”. Journal of Microscopy 195, 10–16. doi: 10.1046/j.1365-2818.1999.00576.x.

26. Marcel Müller et al. (2016). “Open-source image reconstruction of super-resolution structured illumination microscopy data in ImageJ”. Nature Communications 7, ncomms 10980. doi: 10.1038/ncomms10980.

