## Supplemental Code File 1 for "Concepts for structured illumination microscopy with extended axial resolution through mirrored illumination": calculate_otf.m

```
%%%%%%%%%%%%%%%%%%%%%%%%%%%%%%%%%%%%%%%%%%%%%%%%%%%%%%%%%%%%%%%%%%%%%%%%
% Calculate a theoretical OTF for a structured illumination microscope %
% James Manton, 2019                                                   %
%%%%%%%%%%%%%%%%%%%%%%%%%%%%%%%%%%%%%%%%%%%%%%%%%%%%%%%%%%%%%%%%%%%%%%%%


%% Simulation parameters
field_size = 512;
numerical_aperture_primary = 1.27;
numerical_aperture_detection = numerical_aperture_primary;
numerical_aperture_secondary = numerical_aperture_primary;
refractive_index = 1.33;
spot_radius = 0.95;
num_orientations = 3;
INTERFEROMETRIC_DETECTION = 1;
SAVE_IMAGES = 1;
SAVE_3D = 1;
root_name = 'i5s';

primary_alpha = asind(numerical_aperture_primary / refractive_index);
secondary_alpha = asind(numerical_aperture_secondary / refractive_index);


%% Coordinates
lim = field_size - (field_size + 1) / 2;
x = -lim:lim;
y = -lim:lim;
z = -lim:lim;
[X, Y, Z] = ndgrid(x, y, z);
R = sqrt(X.^2 + Y.^2 + Z.^2);
phi = atan2(X, Y) * 180 / pi;
theta = atan2(X, Z) * 180 / pi;
R = 2 * R / max(x);
r = sqrt(X.^2 + Y.^2);
r = 2 * r / max(x);


%% Detection OTF
detection_pupil = zeros(field_size, field_size, field_size);
detection_pupil(r < 0.5 * numerical_aperture_detection / refractive_index) = 1;
detection_ctf = zeros(field_size, field_size, field_size);
if INTERFEROMETRIC_DETECTION
    detection_ctf(R < 0.5 & R > 0.49 & detection_pupil > 0) = 1;
else
    detection_ctf(R < 0.5 & R > 0.49 & detection_pupil > 0 & Z > 0) = 1;
end
detection_otf = fftshift(ifftn(fftn(detection_ctf) .* conj(fftn(detection_ctf))));


%% Primary OTF
primary_pupil = zeros(field_size, field_size, field_size);
primary_pupil(floor(abs(Y)) == round(max(x) * spot_radius * 0.25 * numerical_aperture_primary / refractive_index) & floor(X) == 0) = 1;
primary_pupil(floor(Y) == 0 & floor(X) == 0) = 1;

primary_ctf = zeros(field_size, field_size, field_size);
primary_ctf(R < 0.5 & R > 0.49 & primary_pupil > 0 & Z > 0) = 1;
% Fix band weights
primary_ctf_sum = repmat(sum(primary_ctf, 3), 1, 1, field_size);
primary_ctf(primary_ctf > 0) = primary_ctf(primary_ctf > 0) ./ primary_ctf_sum(primary_ctf > 0);
primary_otf = fftshift(ifftn(fftn(primary_ctf) .* conj(fftn(primary_ctf))));


%% Secondary OTF
secondary_pupil = primary_pupil;

secondary_ctf = zeros(field_size, field_size, field_size);
secondary_ctf(R < 0.5 & R > 0.49 & secondary_pupil > 0 & Z < 0) = 1;
% Fix band weights
secondary_ctf_sum = repmat(sum(secondary_ctf, 3), 1, 1, field_size);
secondary_ctf(secondary_ctf > 0) = secondary_ctf(secondary_ctf > 0) ./ secondary_ctf_sum(secondary_ctf > 0);
secondary_otf = fftshift(ifftn(fftn(secondary_ctf) .* conj(fftn(secondary_ctf))));


%% Dual OTF
dual_ctf = primary_ctf + secondary_ctf;
dual_otf = fftshift(ifftn(fftn(dual_ctf) .* conj(fftn(dual_ctf))));


%% Overall OTF
detection_psf = otf2psf(detection_otf);
dual_psf = otf2psf(dual_otf);
overall_psf = detection_psf .* dual_psf;
overall_otf = fftshift(psf2otf(overall_psf));

all_rotations_otf = overall_otf;
all_rotations_otf = all_rotations_otf ./ max(all_rotations_otf(:));
if (num_orientations > 1)
    for i = 2:num_orientations
        rotated_otf = imrotate(overall_otf, (i - 1) * 180 / num_orientations, 'bilinear', 'crop');
        rotated_otf = rotated_otf ./ max(rotated_otf(:));
        all_rotations_otf = all_rotations_otf + rotated_otf;
    end
end
overall_otf = all_rotations_otf;
overall_psf = real(otf2psf(fftshift(overall_otf)));


%% Display
primary_ctf_pxz = squeeze(sum(primary_ctf, 1))';
primary_ctf_pyz = squeeze(sum(primary_ctf, 2))';
primary_ctf_pxy = squeeze(sum(primary_ctf, 3));

secondary_ctf_pxz = squeeze(sum(secondary_ctf, 1))';
secondary_ctf_pyz = squeeze(sum(secondary_ctf, 2))';
secondary_ctf_pxy = squeeze(sum(secondary_ctf, 3));

dual_otf_pxz = log(1 + squeeze(sum(dual_otf, 1))');
dual_otf_pyz = log(1 + squeeze(sum(dual_otf, 2))');
dual_otf_pxy = log(1 + squeeze(sum(dual_otf, 3)));

overall_otf_pxz = (squeeze(sum(overall_otf, 1))');
overall_otf_pyz = (squeeze(sum(overall_otf, 2))');
overall_otf_pxy = (squeeze(sum(overall_otf, 3)));

overall_otf_sxz = (squeeze(overall_otf(:, field_size / 2, :)))';
overall_otf_syz = (squeeze(overall_otf(field_size / 2, :, :)))';
overall_otf_sxy = (squeeze(overall_otf(:, :, field_size / 2)));

overall_psf_pxz = (squeeze(sum(overall_psf, 1))');
overall_psf_pyz = (squeeze(sum(overall_psf, 2))');
overall_psf_pxy = (squeeze(sum(overall_psf, 3)));

overall_psf_sxz = (squeeze(overall_psf(:, field_size / 2, :)))';
overall_psf_syz = (squeeze(overall_psf(field_size / 2, :, :)))';
overall_psf_sxy = (squeeze(overall_psf(:, :, field_size / 2)));

figure(1)
imshow([primary_ctf_pxz ./ max(primary_ctf_pxz(:)), primary_ctf_pyz ./ max(primary_ctf_pyz(:)), primary_ctf_pxy ./ max(primary_ctf_pxy(:))], [])
title('Primary CTF projection')

figure(2)
imshow([secondary_ctf_pxz ./ max(secondary_ctf_pxz(:)), secondary_ctf_pyz ./ max(secondary_ctf_pyz(:)), secondary_ctf_pxy ./ max(secondary_ctf_pxy(:))], [])
title('Secondary CTF projection')

figure(3)
imshow([dual_otf_pxz ./ max(dual_otf_pxz(:)), dual_otf_pyz ./ max(dual_otf_pyz(:)), dual_otf_pxy ./ max(dual_otf_pxy(:))], [])
title('Dual OTF projection')

figure(4)
overall_proj_display = [overall_otf_pxz ./ max(overall_otf_pxz(:)), overall_otf_pyz ./ max(overall_otf_pyz(:)), overall_otf_pxy ./ max(overall_otf_pxy(:))];
imshow(overall_proj_display, [])
title('Overall OTF projection')

figure(5)
overall_slice_display = [overall_otf_sxz ./ max(overall_otf_sxz(:)), overall_otf_syz ./ max(overall_otf_syz(:)), overall_otf_sxy ./ max(overall_otf_sxy(:))];
imshow(overall_slice_display, [])
title('Overall OTF slice')

figure(6)
overall_psf_proj_display = [overall_psf_pxz ./ max(overall_psf_pxz(:)), overall_psf_pyz ./ max(overall_psf_pyz(:)), overall_psf_pxy ./ max(overall_psf_pxy(:))];
imshow(overall_psf_proj_display, [])
title('PSF projection')

figure(7)
overall_psf_slice_display = [overall_psf_sxz ./ max(overall_psf_sxz(:)), overall_psf_syz ./ max(overall_psf_syz(:)), overall_psf_sxy ./ max(overall_psf_sxy(:))];
imshow(overall_psf_slice_display, [])
title('PSF slice')


if (SAVE_IMAGES)
    imwrite(uint16((2^16-1) * overall_proj_display), [root_name, '_otf_', num2str(num_orientations), 'a_', strrep(num2str(numerical_aperture_primary), '.', 'p'), '_', strrep(num2str(numerical_aperture_secondary), '.', 'p'), '_p.tif'])
    imwrite(uint16((2^16-1) * overall_slice_display), [root_name, '_otf_', num2str(num_orientations), 'a_', strrep(num2str(numerical_aperture_primary), '.', 'p'), '_', strrep(num2str(numerical_aperture_secondary), '.', 'p'), '_s.tif'])
    imwrite(uint16((2^16-1) * overall_psf_proj_display), [root_name, '_psf_', num2str(num_orientations), 'a_', strrep(num2str(numerical_aperture_primary), '.', 'p'), '_', strrep(num2str(numerical_aperture_secondary), '.', 'p'), '_p.tif'])
    imwrite(uint16((2^16-1) * overall_psf_slice_display), [root_name, '_psf_', num2str(num_orientations), 'a_', strrep(num2str(numerical_aperture_primary), '.', 'p'), '_', strrep(num2str(numerical_aperture_secondary), '.', 'p'), '_s.tif'])
end

if (SAVE_3D)
   otf_16 = uint16((2^16 - 1) * overall_otf ./ max(overall_otf(:)));
   imwrite(otf_16(:, :, 1), [root_name, '_otf_', num2str(num_orientations), 'a_', strrep(num2str(numerical_aperture_primary), '.', 'p'), '_', strrep(num2str(numerical_aperture_secondary), '.', 'p'), '_3D.tif'])
   for i = 2:size(otf_16, 3)
      imwrite(otf_16(:, :, i), [root_name, '_otf_', num2str(num_orientations), 'a_', strrep(num2str(numerical_aperture_primary), '.', 'p'), '_', strrep(num2str(numerical_aperture_secondary), '.', 'p'), '_3D.tif'], 'WriteMode', 'Append') 
   end
end
```
